# A Mathematical model to predict Trichodesmiun *spp*. bloom during a sewage burst

**DOI:** 10.1101/179853

**Authors:** Guido Santos Rosales

## Abstract

A modified version of the Monod equation is proposed to understand the bloom emergence of *Trichodesmiun spp*. in presence of sewage outburst. The mathematical model predicts that the bacterial population does not affect the bloom growth. Also, the quotient between the bacteria during the sewage outburst and before is proposed to be equal to the quotient between the influx of limiting resources during burst and the influx before the burst.

## Introduction

*Trichodesmiun spp*. is a N_2_-fixing cyanobacterium responsible for bacterial blooms^1^. The emergence of these blooms depends on temperature, local marine currents and micronutrients like Fe^2^. Recently is has been stablished a relationship between sewage bursts and *Trichodesmiun* blooms^3^. In order to understand the quantitative evolution of these bursts and the effect of sewage influx here it is proposed a mathematical quantification of the process using quantitative data from the previous study.

## Methods

*Experimental data.* The data was taken from a study about a *Trichodesmiun spp*. bloom^3^. The bloom was followed before, during and after a sewage burst in the south-eastern Mediterranean Sea. The authors measured a time-series concentration of *Trichodesmiun spp*. cells in the region where the sewage outburst happened. Figure 1 displays the data.

**Figure 1.**
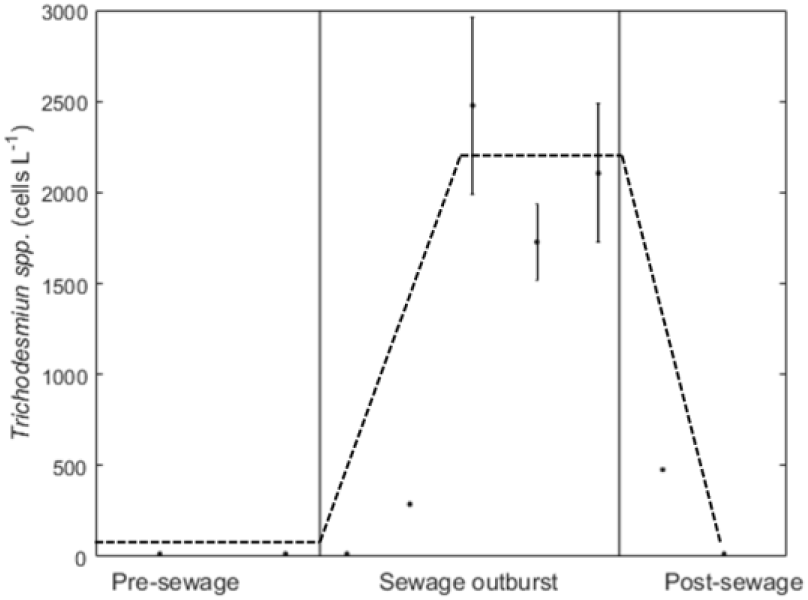
Experimental data from Rahav & Bar-Zeev 2017^3^. Dots represent the data, bars represent the experimental error when available and dashed lines show the qualitative pattern that it is pretended to be predicted.

*Parameter estimation*. The different models were fitted to the data with a simple estimator using the squared error to the data as objective. After an initial set of random values between 0 and 1 for the parameters of the model the next set of parameters is set randomly using a normal distribution centred in the previous value with a standard deviation of a tenth of this value. This estimator is able to follow the minimal in an unbounded way. For each model the estimator is run during 10^4^ iterations, and the best solution is saved. 10 solutions for each model are analysed.

## Results

The experimental data^3^ show the evolution of a population of *Trichodesmiun spp*. before, during and after a sewage outburst. In the data it is observed a steady phase during the sewage outburst (Figure 1). Here, it is going to be assumed that the bacteria stop growing not because the nutrients are exhausted. This hypothesis is supported by the fact that the outburst was continued during this period.

The well stablished mono-substrate Monod equation^4^ (*Model 0*) is used as a starting point to predict the bacterial growth. *R* represents the limiting resource for growing and *B* the bacteria count.

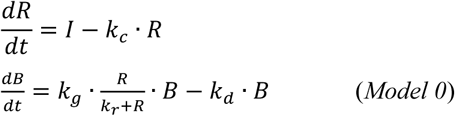

The steady phase of the bacteria during the sewage cannot be explained by the Monod equation, because the steady state in this model is only satisfied for a restricted relationship of the parameters. And the value of the steady state does not depend on the values of the parameters. To solve this we assume that bacteria size is not a limiting factor for growth (*Model 1*). One way to see this is assuming that it is a restriction in the environment that constrains the maximal bacterial production rate, like a carbon resource or minerals. Here it is assumed that the resource is the limiting necessity for growing that can be a carbon resource, a mineral component or another environmental condition that restricts the growth. The only assumption is that this resource is increased during the sewage outburst (*I*^*Sewage*^ > *I*^*Pre-Sewage*^).

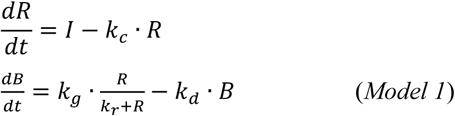

Monod equation assumes saturation in the income of the resource. But this assumption introduces an additional parameter in the model. For the lack of data we are going to consider the lineal model too (*Model 2*).

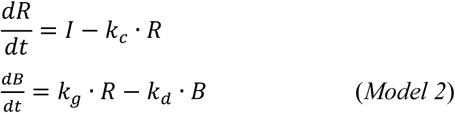

Figure 2 shows the fit of *Model 1* and *Model 2* to the data. *Model 1* fits the maximal data better than *Model 2*, but *Model 2* presents a better fit for the data post-sewage.

**Figure 2.**
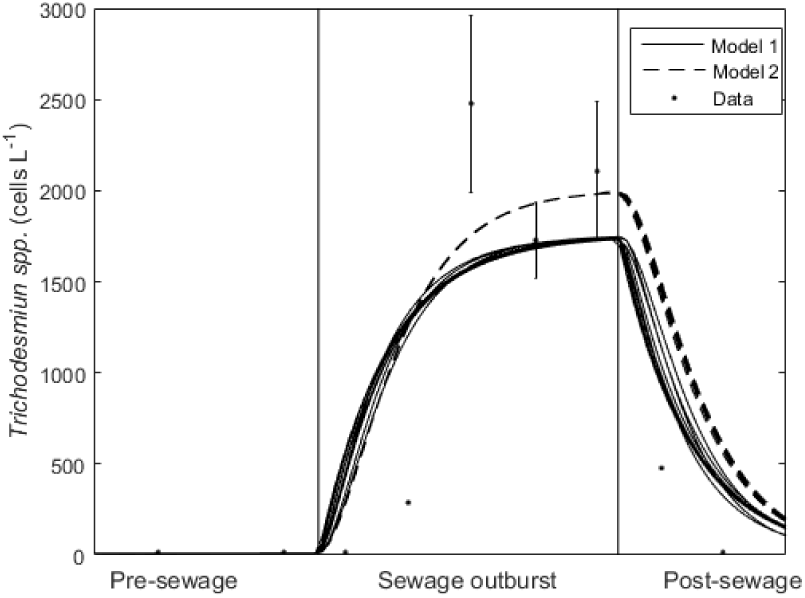
*Model 1* and *Model 2* fitted to experimental data from Rahav & Bar-Zeev 2017^3^. 10 solutions for each model were plotted.

*Model 2* has the best fit to the whole set of data according to the squared error. Table 1 contains the detailed information of the fit errors. It is interesting that *Model 2* presents a better fit even with one parameter less than *Model 1*.

**Table 1.** Squared errors of the fit between models to data. Mean and standard deviation (s.d.) of 10 solutions.

|  | Parameters | mean | s.d. |
| --- | --- | --- | --- |
| *Model 1* | 4 | $2.51 \cdot 10^6$ | $9.40 \cdot 10^4$ |
| *Model 2* | 3 | $2.30 \cdot 10^6$ | $8.01 \cdot 10^3$ |
| *Model 3* | 5 | $1.43 \cdot 10^6$ | $5.97 \cdot 10^4$ |

One way to improve the fit without increasing too much the complexity of the model is working with the generalized version for a higher number of resources. *Model 3* introduces additional resources in the equation that can affect the bacterial growth.

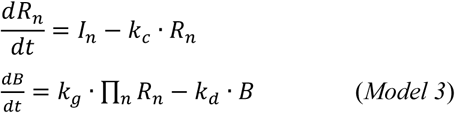

In the case of two resources the model improves the fit as it can be seen in Figure 3. *Model 3* is able to fit as good as *Model 2* the higher values of bacteria, and at the same time it reduces the error in the data post-sewage.

**Figure 3.**
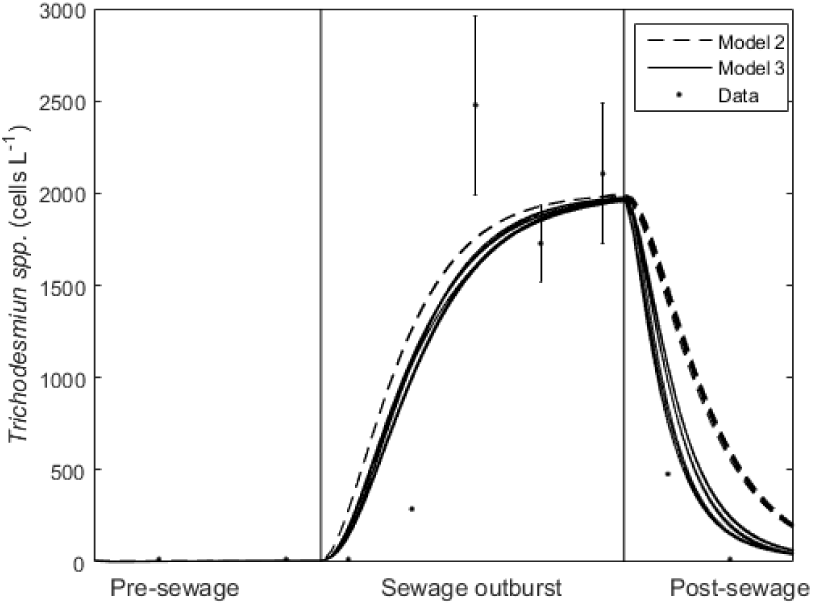
*Model 2* and *Model 3* fitted to experimental data from Rahav & Bar-Zeev 2017^3^. For *Model 3* it was used 2 resources. 10 solutions for each model were plotted.

Table 1 shows that the global squared error is much lower in *Model 3* than in *Model 2*. We should take into account that this model has 2 more parameters, which can facilitate the fit. But we decided for the multi-resource model because the increase in complexity on the model is slight compared with other strategies and it is natural to assume that the bacteria will have more than one restricting resource in these conditions.

The quotient of the change between steady state of bacteria during the sewage outburst and before the burst, given by *Model 3*, is equal to

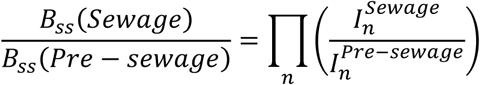

where 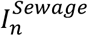 is the influx of the resource *n* during the sewage and 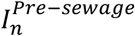 is the influx of the resource *n* in natural conditions, before the sewage burst started.

## Discussion

In this work it is assumed a simple mathematical model to predict a bloom of *Trichodesmiun spp*. after a sewage outburst. The data present a singular pattern that cannot be predicted using the Monod equation for bacterial growth. This model can be improved if it is assumed that bacteria growth at a rate proportionally to the limiting resource that can be any kind of resource, not only a carbon source. It should be considered that the maximal growth of biomass is limited by the availability of certain resource.

A simpler version of the model, assuming a linear relationship between the resource and the growth is able to fit the data better than the model with saturation in the resource intake, even when the linear model has a lower number of parameters. This observation is suggesting that the limiting resource for the bloom may be consumed without saturation. This could be because the availability of the limiting resource is low or because the bacterial uptake machinery for this resource is very efficient.

The model can be further improved if we assume that there is more than one limiting resource. When we assume that there are two limiting resources the fit to the data is better than modelling only one limiting resource. This explanation is natural because *Trichodesmiun spp*. could have been behaving as mixotrophic during the bloom emergence to deal with the high energetic cost of bloom formation.

Assuming the previous model, we can calculate the change of *Trichodesmiun spp*. during the bloom. The quotient between the bacteria during the sewage outburst and before the outburst is equal to the product of the quotients between the influx of resource during burst and the influx in natural conditions.

The present work is just a theoretical approach to the problem of cyanobacterial blooms and the possible effect of sewage burst. Here it is used real data to calibrate the models that were used to make predictions. Although the data in insufficient to obtain accurate predictions some interesting quantitative relationships can be inferred. This work demonstrates the utility of quantitative analysis to understand complex problems like bacterial blooms. Following the blooms with systematic measurements and combining these data with quantitative analysis would provide a better understanding to the problem and it could propose well informed strategies to control their emergence.

